# Base composition at the start of the coding sequence controls the balance between translation initiation and mRNA degradation in *E. coli*

**DOI:** 10.1101/2024.03.21.586065

**Authors:** Anna Lipońska, Laura Monlezun, Isaac Wilkins, Saravuth Ngo, Thomas Oïffer, Cylia Bouchachi, John F. Hunt, Daniel P. Aalberts, Grégory Boël

## Abstract

Protein synthesis efficiency is highly dependent on the mRNA coding sequence. Furthermore, there is extensive evidence of a correlation between mRNA stability and protein expression level, though the mechanistic determinants remain unclear. Using yellow fluorescent protein (YFP) as a reporter gene, we herein demonstrate that adenosine (A) abundance in the first six codons is a critical determinant for achieving high protein synthesis in *E. coli*. Increasing A and/or decreasing guanosine (G) content in this region with synonymous codons results in substantial increases in protein expression level both *in vivo* and *in vitro* that are correlated with steady-state mRNA concentration *in vivo*. The change in mRNA concentration is attributable to changes in the stability of the mRNA that are directly coupled to its translation efficiency. Increasing A content promotes mRNA incorporation into the functional 70S ribosomal initiation complex without altering its affinity for the 30S ribosomal subunit. These results support a model in which base composition in the first six codons modulates local mRNA folding energy and single-strandedness to control the balance between productive translation initiation *versus* degradation of mRNAs bound to the 30S ribosomal subunit. Based on these findings, we developed a short N-terminal coding sequence that optimizes translation initiation efficiency for protein production in *E. coli*.

## Introduction

Biological protein synthesis comprises the translation of mRNA by the ribosome into protein. Since the genetic code uses 61 nucleotide codons to encode the 20 amino acids (aa) contained in proteins, there is a degeneracy that allows a large number of mRNA sequences using synonymous codons to encode the same protein sequence. Nevertheless, synonymous sequences (sequences which encode for the same aa but use different codons) can strongly affect protein expression (1-8). The mechanisms behind this effect remain unclear in part because synonymous codons can influence translation in different ways. First, they change the base composition and therefore the structure of the mRNA. Secondly, they can use different tRNA with different abundance or affinity.

Several studies have shown that codons can change translation efficiency during elongation of the nascent protein (4,9-16). It has been widely believed that tRNA abundance, which is correlated with *Escherichia coli* codon usage (6,10,17), controls translation elongation efficiency (1,4,6,7,9,10,18-20). This model postulates that the so-called rare codons can slow ribosome elongation, but recent publications have challenged this view (11,13,15,16,21-33). Ribosome profiling and high throughput protein expression experiments have established that translation elongation efficiency can vary, but there is only a weak correlation with tRNA abundance (13,15,16,23,34-36). Moreover, these studies have shown that codons influence mRNA translation and that there is a correlation with the translation of mRNA and mRNA stability (13,37-41), possibly related to ribosomal stalling and collision (42,43).

Rare codons become problematic only when they are overused, for example, when multiple repeats of a rare codon are inserted into a sequence or when a gene with a high frequency of rare codons is strongly overexpressed (4,7). This effect arises from depletion of the pool of charged (aminoacylated) tRNAs, not from a lower overall abundance of the corresponding tRNA species (14,44).

Varying results have been obtained from experiments in *E. coli* using libraries of fluorescent reporter genes in which the first codons are completely randomized (45,46) or substituted by synonymous codons (11,21). First, the calculated free energy of mRNA folding in the 5’UTR and the initial segment of the coding sequence correlates with expression level, with less stable folding (higher ΔG) promoting increased protein expression (11,21,45,46). An equivalent correlation was also observed in a study where all the codons of a fluorescent protein were randomized at synonymous sites (47). This study also observed an A bias in high-expressing constructs (codons 2–7), whereas G at codon 4 and C at codons 2, 3, and 7 were negatively associated with expression. Second, the base composition in the initial codons shows increased expression with a higher frequency of Adenosine (A) (45,46). A study by Goodman *et al*. shows that use of the rarest codons in this region favor higher expression (11). These codons are enriched in A/U compared to G/C (17,46), which are likely responsible for beneficial effect base on statistical analyses large-scale expression experiments (13). The Goodman *et al*. paper (11) furthermore demonstrate that the abundance of tRNA cognate to codons in this region is not a significant determinant of expression.

In a study focusing on codons 3-5 of the coding sequence, Verma e*t al*. reported data supporting specific amino acid motifs at these sites can influence protein expression (45). The largest effects were found for nucleotide base patterns AADUAU (where D stands for any base except C) and AAVAUU (where V stands for any base except U). These mRNA sequences correspond to protein sequences with Lys or Asp at position 3 and Tyr or Iso at position 4 (K/N-Y/I). The authors propose that the protein-expression-enhancing effects of these sequence derive in part from reductions in ribosome pausing and premature termination of protein synthesis at codons 3-5 (45). Indeed, during the first rounds of elongation after initiation, the peptidyl-tRNAs can dissociate from the ribosome when problematic amino acids are incorporated into the nascent polypeptide (45,48-50). This process, known as peptidyl-tRNA drop-off, is proposed to be, in part, a mechanism by which the ribosome rejects miscoded peptidyl-tRNAs during early elongation (48).

However, for most mRNA, the rate-limiting step of protein synthesis is the formation of a 70S elongation-competent complex (70S EC), a mature form of the 70S initiation complex (70S IC) (51-53). The rate of 70S EC formation depends on the mRNA and ranges between 0.5 sec to a few seconds (52,54,55), while translation elongation occurs faster at an average speed of ∼20 amino acids per second (56,57). The current general model of 70S IC assembly starts with the formation of the 30S pre-IC, which is formed by the three initiation factors (IF1, 2 and 3), the initiator tRNA (fMet-tRNA^fMet^) and the 30S ribosomal subunit (58). This complex then associates with the mRNA at the initiation codon, which, in most cases in *E. coli*, is guided by a Shine-Dalgarno sequence (SD) upstream of this codon and by the nucleotide composition of the surrounding region (17,59). Finally, joining of the 50S ribosomal subunit to the 30S pre-IC triggers IF-3 dissociation to form the 70S pre-IC, followed by repositioning of fMet-tRNA^fMet^ and departure of IF1 to create the 70S IC. Subsequent dissociation of IF2 generates the 70S EC (52,58). The kinetics of each of these steps determines the effective translation initiation rate (TIR).

TIR also determines the density of ribosomes on translating mRNAs, thus influencing ribosome collision frequency (60). It has been proposed that the collision of a stalled ribosome by a trailing ribosome stimulates abortive termination of the stalled ribosome, and this effect is dependent on the TIR (61). The TIR can be modulated by the folding of the sequence upstream of the initiating codon (5’UTR) and the initiating codon (21,34,62-65). Furthermore, mRNA folding extending 40 nucleotides into the coding sequence can also influence TIR (13,21). The mRNA secondary structures formed in this region can be modified by environmental conditions such as temperature or interaction with *trans*-acting regulators, small RNA, or ribosomes to regulate protein expression (66). The SD can also modulate the formation of the initiation complex by its complementarity with the anti-SD sequence of the 16S rRNA and its position in the sequence (67-70).

In the region of the mRNA surrounding the ATG, several studies have shown that nucleotide base biases toward adenosine (A) correlates with an increase in protein expression (59,71,72). We additionally found that guanosine (G) in the first 18 nucleotides of the coding sequence reduces protein expression (13,73). Bias in nucleotide base composition is clearly observed in this region of protein coding sequences in the *E. coli* K12 genome (**Supplementary Figure S1**). This small region has been proposed to control binding to the 30S pre-IC via ribosomal protein S1 (74,75) to form the 30S IC, which is an essential precursor to formation of the functional 70S IC. Furthermore, this mRNA region has furthermore been implicated in guiding alignment of the ribosome at the proper initiation codon (59).

In the work presented here, we employ a Yellow Fluorescence Protein (YFP) reporter gene to characterize the influence of the nucleotide base composition in the first six codons. We modify the A and G fractions of the first six codons identified in our previous work (13) and included an additional codon to slightly extend the original window. These changes were introduced either by synonymous codon substitutions or by appending the reporter to the first 21 nucleotides of a highly expressed endogenous *E. coli* gene. Fluorescence measurements show that increasing A content and/or decreasing G content produces up to 10-fold increases in expression of the YFP reporter. In all the reporters studied, improvement in protein expression results in an increase of the mRNA stability. Furthermore, we demonstrate that the 5’ coding sequence that produces the greatest increase in YFP expression, which we named “Translation Boosting Tag” (TBT), also strongly enhances expression of poorly expressed proteins when fused in front of their coding sequences in a T7 expression system. Finally, we present biochemical studies demonstrating that modulating A and G content in the first six codons at the 5’ end of the coding sequence does not change mRNA affinity for the 30S ribosomal subunit, but it does change the efficiency with which that complex progresses to form a productive 70S IC. These results suggest that the formation of the 70S IC from the 30S IC controls a competition between translation initiation and mRNA decay that coordinates these two processes based on the nucleotide base content in the ribosome-docking region at the start of the coding sequence. We propose that this mechanism is a central feature of mRNA metabolism in *E. coli* that optimizes the use of metabolic resources by initiating mRNA decay upstream of translating ribosomes rather than downstream where decay would lead to wasting the energy used to synthesize incomplete proteins.

## Materials and Methods

### Plasmids and strains

*E. coli* strain DH5α (76) was used for plasmid constructions. All experiments with a use of pMMB67EH plasmids (77) were performed in the MG1655 *E. coli* strain (78). Proteins overexpression with pET21 plasmids were carried out in *E. coli* strain BL21 (DE3) (79). Cultures were grown at 37 °C in Luria-Bertani Miller broth (LB-Difco) supplemented with ampicillin at 100 µg/ml. Plasmids were constructed by PCR amplification of whole plasmids with the CloneAmp Hifi polymerase (Takara) and assembly of the insert part with the NEBuilder HiFi DNA Assembly kit. The *yfp* variants plasmids were constructed by amplification of the pMMB67EH (77) backbone with primers and amplification of the *yfp* gene (80,81) with primers that contain or lack the modification of the first codon of the *yfp* or that add the different TBT sequences; the primers are described in **Table 1**. The same approach was used to insert the TBT sequence in the pET21 expressed test genes *ycaQ*, RSP_2139, SCO1897 and SRU_1983 that we have used in our previous study (13). A 38nt RNA malachite green aptamer (MGapt) was added as described by Siegal-Gaskins *et al*., (82) in the 3’UTR of the reporter mRNA, 10nt downstream of the stop codon of the TBT or TBT_G_ *yfp* genes. To that end, a PCR amplification of whole plasmids was performed with primers H070 and H071, which contain 20 bases of homology to allow plasmid recircularization using the NEBuilder HiFi DNA Assembly kit.

**Table 1:**
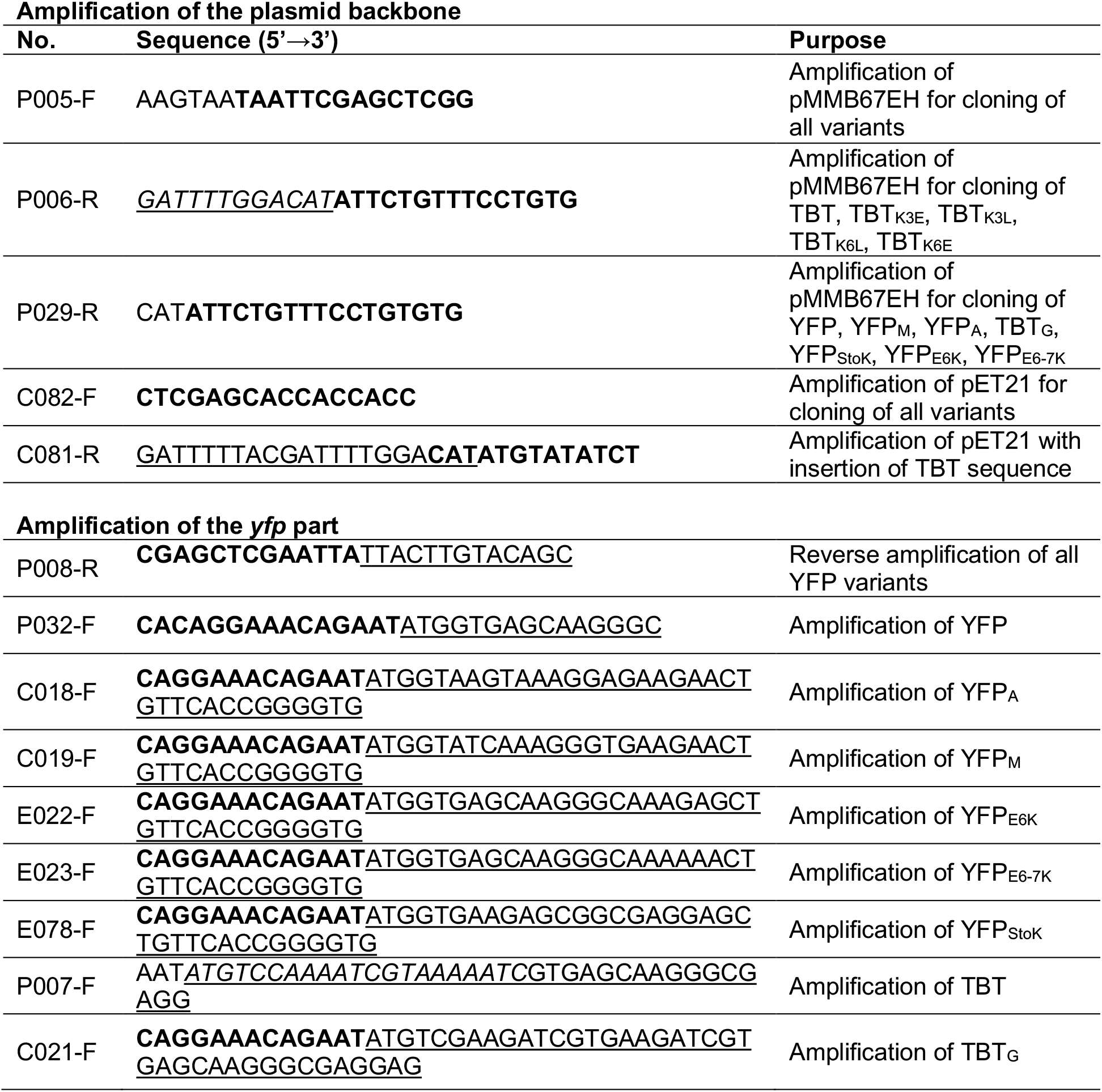
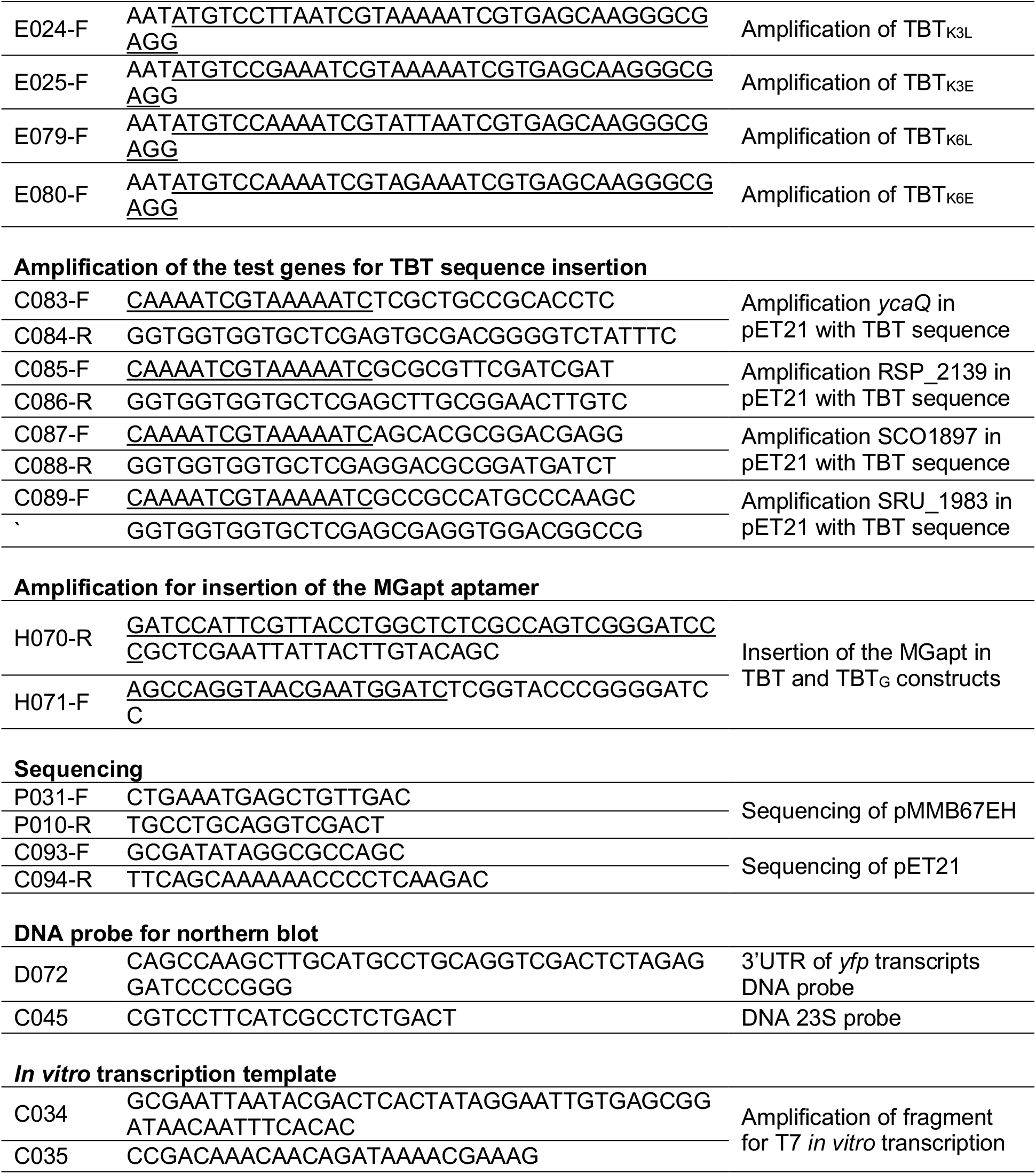
Oligonucleotides. Sequences in bold are homologous to the plasmid backbone, while sequences underlined contain the modifications in the first codon of the *yfp* or the MGapt aptamer.

### Drop growth assay

Samples corresponding to OD_600_=1 of bacterial overnight cultures were centrifuged and diluted in 1 ml LB from which further dilutions were prepared: 10^-1^, 10^-2^, 10^-3^, 10^-4^, 10^-5^. 5 µl of sample was placed on LB ampicillin (100 µg/ml) plate and grown overnight (O/N) at 37 °C. Fluorescence was measured using a Typhoon FLA 9 500 apparatus at 473 nm, using the BPB1 filter. The results for the best dilution are presented in the figures. Experiments were repeated 3 times.

### *In vivo* measures of YFP and TBT variants

Bacteria were grown in Falcon 96-well sterile plates in the CLARIOstar (BMG LabTech) device to quantify optic density of bacteria and the emission of YFP fluorescence (ex 497-15/em 540-20 gain 1000, measured from top 4.0 diameter every 30 min). 2 µl of overnight culture was added directly to 200μl of LB ampicillin (100 µg/ml) (without or with 1 mM IPTG), 60μl of mineral oil (Sigma Life Science) was added to prevent evaporation of the medium and to keep sterile conditions. Each culture was made in triplicate. Data were analyzed with Prism 8 software.

For the experiments with sample collection for northern and western blot, bacteria were grown at 37 °C until OD_600_=0.5, then 1 mM IPTG was added. The samples were collected at 0, 10, 60 and 120 min. For protein samples an equivalent sample for 1 OD was collected and centrifuged, the pellets were resuspended in 200 µl of 1x Laemmli sample buffer (Bio-Rad) and boiled for 2 min at 95°C before migration on SDS denaturing gel. For the RNA sample, 1 ml of culture were collected and centrifuged, resuspended in 150 µl of RNAsnap (83) and processed as described for the RNAsnap procedure. Samples were analyzed on a 12.5% SDS-PAGE gel, by western blot and on agarose gel by northern blot.

### Test genes

Plasmids encoding genes *ycaQ* (*E. coli*), RSP_2139 (*Rhodobacter sphaeroides*), SCO1897 (*Streptomyces coelicolor*) and SRU_1983 (*Salinibacter ruber*) were previously assessed for their expression in the *E. coli* Bl21 DE3 strain (13). We introduced the TBT sequence in the 5’-end of the coding sequence (see above). Then all the constructs with or without the TBT sequence were grown at 37 °C to OD_600_=0.5 in LB with Amp 100 µg/ml. Protein expression was induced by addition of 1 mM IPTG for 16h at 17 °C. Samples of 1 OD were collected before and after induction and analyzed on a 12.5% SDS-PAGE gel and by western blot.

### Western Blot

Equal amounts of Total Protein extracts were migrated on a 12.5% SDS-PAGE gel and transferred onto Immun-Blot PVDF blotting membrane (Bio-Rad). Transfer was performed with the Bio-Rad Transfer System (30 min, 25V). For YFP detection, an anti-GFP polyclonal antibody was used (Invitrogen, 11122) in dilution 1:1000 in 1X PBS, 0.1 % Tween-20 and detected with an anti-rabbit IRdye 800CW (LI-COR, 926-32213) in dilution 1:20,000 in the same buffer. For proteins with His-tag, we used anti-6xHis-tag primary antibody (Covalab, ref. HIS.HS/ EH158) in dilution 1:2,000 detected with anti-mouse IRdye 800CW (LI-COR, 926-32210) in dilution 1:20,000. The membrane was visualized with Odyssey scanner (LI-COR).

### mRNA stability assay

Bacteria were grown until OD_600_= 0.5, then 1 mM IPTG was added. Induction was carried out for 10 min and was stopped by adding of rifampicin (0.5 mg/ml; 0.6 mM). Samples were collected as indicated after 1, 2, 3 and 4 minutes; to stop cell growth 10 mM sodium azide was added. The mixture was vortexed and centrifuged (1 min/ RT/ 16 000 x g) and RNA were isolated with the RNAsnap method(83).

### *In vitro* transcription

The DNA templates used to generate mRNAs for *in vitro* translation assays were PCR amplified with primers C034 and C035 from the corresponding pMMB67EH plasmids. The forward primer contains the T7 polymerase promoter sequence (5’-GCGAATTAATACGACTCACTATAGGG-3’). *In vitro* mRNA synthesis was then carried out at 37 °C for 4 h in T7 the RiboMAX Large Scale RNA Production System kit (Promega) according to the manufacturer’s recommendations. At the end of the reaction, DNA templates were degraded by adding DNAse I for 15 min and the mRNA transcripts were purified using TRIzol Reagent (Thermo Fisher Scientific) and Direct-zol RNA Miniprep kit (Zymo Research). The mRNAs were finally eluted in The RNA Storage Solution (Ambion, Thermo Fisher Scientific) and stored at −80 °C.

### *In vitro* translation

To test the expression of the different constructs, 1.4 µM of each purified mRNA was used in the PURExpress *in vitro* Protein Synthesis Kit (NEB#E6800) in a 10 µl reaction in a 384 well Small Volume black plate (Grenier 788 096). Fluorescence was measured with CLARIOstar (BMG LabTech; ex 497-15/em 540-20 gain 1000, measure every 4 min) during 4 hours at 37 °C. After reaction, 2 µl of reaction was stopped with 4x Laemmli sample buffer (Bio-Rad) and used for western blot analysis. For the IVTA separated on sucrose gradient (**Figure 4B**) reactions were performed in a final volume of 100 µl using the PURExpress ΔRibosome *In vitro* Protein Synthesis Kit (New England Biolabs) according to the manufacturer instructions and with ribosomes purified from *E. coli* strain MRE600(49) at a final concentration of 2 μM. The reporter mRNA was added at a final concentration of 1 µM and the reaction was then carried out at 37 °C for 1h. The translation of the reporter mRNA was monitored every 2 min by measuring YFP fluorescence on a CLARIOstar plate reader (BMG Labtech).

### Polysomes fractionation on sucrose gradient and RNA precipitation

Entire *in vitro* translation reactions were loaded onto 10-40 % (w/v) sucrose gradients (20 mM Tris pH 8, 20 mM MgCl_2_, 100 mM NH_4_(OAc), 2 mM β-mercaptoethanol) and centrifuged 2 h at 200,000 g, 4 °C in SW41 rotor (Beckman Coulter). Polysomes profiles were obtained using Biocomp Piston Gradient Fractionator (BioComp Instruments) by measuring absorbance at 254 nm, and 400 μl fractions were collected in 96 wells plates (Thermo Scientific).

Fractions collected from the polysomes profiles were precipitated with phenol/chloroform. One volume of phenol/chloroform/isoamyl alcohol was added to each fraction and samples were centrifuged at 18,000 g for 5 minutes at 4 °C. The aqueous upper phase was carefully collected. Sodium acetate 3M (1/10 of sample volume), pure ethanol (2,5 volume of sample volume) and Glycogen were added to the collected phase, then samples were incubated overnight at −80 °C. Samples were centrifuged at 18,000 g for 30 minutes at 4 °C, and the supernatants were then discarded. Pellets were washed with 200 μl of ethanol 70% (v/v) and centrifuged again at 15,000 rpm for 15 minutes at 4 °C. Pellets were set to dry on the bench before being resuspended in 20 μl of The RNA Storage Solution (Ambion, Thermo Fisher Scientific) and stored at −80 °C.

### Northern Blotting

Precipitated RNAs (5 μl) or RNAsnap extracts were loaded on a 1% (w/v) agarose gel and then transferred to an Amersham Hybond-N+ membrane (Cytiva). After migration, 16S and 23S RNA were visualized with Ethidium Bromide (EtBr) under UV. RNAs were cross-linked on the membrane by exposing the membrane to UV for 30 s at 100 x 1 200 µJ / cm^2^. The radioactive probe was prepared from 40 pmoles of D072 primer (**Table 1**) and labeled at the 5’ end with 10 units of T4 polynucleotide kinase (New England Biolabs) and [*γ*-^32^P]-ATP (150 µCi). After pre-incubation for 1 hour at 42 °C in the hybridization buffer (ULTRAhyb, Invitrogen Thermo Fisher Scientific), membranes were incubated with the probe, in the hybridization buffer, overnight at 42 °C. Membranes were washed three times (once in 2x SSC + 0.1% (v/v) SDS, once in 1x SSC + 0.1% (v/v) SDS and finally in 0.1x SSC + 0.1% (v/v) SDS) at 42 °C during 10 min. Radioactive signal was detected on a Typhoon 9 500 FLA scanner (GE Healthcare). Quantification of the EtBr and radioactive signals was performed with Fiji software (84). For the mRNA quantification experiments a loading control was done by rehybridizing the membrane with the 23S rRNA probe (C045, **table 1**).

### Peptidyl-tRNA drop off assay

Experiments were performed as described by Chadani *et al*. (50). with minor modifications. Coupled transcription-translation reactions were carried out using PUREfrex 1.0 (GeneFrontier) at 37°C for 90 min, containing [^35^S]-methionine (0.2 µCi/µl), 2 µM purified ribosomes from *E. coli* MRE600 (49) and 4 ng DNA templates for *in vitro* translation assays (see above). Reactions were stopped with 1 mL of 5% TCA, incubated on ice for 10 min, centrifuged (3 min, 4°C), and pellets washed with 0.9 mL acetone. Precipitates were dissolved in SDS sample buffer (62.5 mM Tris-HCl pH 6.8, 2% SDS, 10% glycerol, 50 mM DTT). Samples were split; one portion treated with 50 μg/mL RNase A (Promega) at 37°C for 30 min then separated by electrophoresis on a 11% WIDE Range Gel (Nacalai Tesque).

### Filter assay

To test the binding affinity of the TBT and TBT_G_ mRNA to the 30S ribosomal subunit we used PURExpress ΔRibosome Kit (NEB) supplemented with 5 µM of 30S ribosomal subunit and 10 µM tRNA^fMet^ in the presence of 1.4 µM of radioactively labelled mRNA (prepared as described above by *in vitro* translation followed by [^32^P] labelling as described for the northern blot probe labelling). 30S ribosome subunit were purified from *E. coli* strain MRE600 (49). The reaction was performed at 4 °C in the buffer: 20 mM Tris-HCl pH 7; 15 mM MgOAc; 100 mM NH_4_OAc; 1 mM DTT. At 30, 60 and 300 sec, 4 µl of the reaction were taken and deposited on a nitrocellulose filter (MF-Millipore 0.025 µm MCE Membrane) right after 2x 6 ml of reaction buffer was used to wash the filter under vacuum. Radioactive signal was detected on a Typhoon 9500 FLA scanner (GE Healthcare) and quantification of the signals was performed with Fiji software (84).

### Computational modeling

In Figure 2, we observe that after transcription is blocked, some portion of the mRNA decays within the first minute, and the remaining mRNA decays exponentially with a measurable rate Γ:

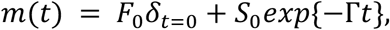

where *δ*_*t*=0_ = 1 when *t=0* and *δ*_*t*=0_ = 0 for *t* ≥1 min. We will now describe how we arrive at this functional form with our proposed fast and slow population model of Figure 3C,

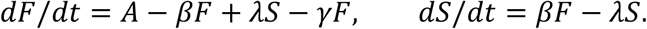

The steady-state values for the Fast and Slow populations are

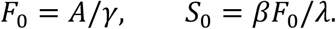

Blocking transcription sends *A* → 0. The mRNA shows a bi-phasic decay, one with a decay time much less than 1 min and the other on the order of few minutes. To match the model, if we take *γ* ≫ 1/min, we obtain the very rapid decay of the *F* population, approximating the delta-function above. Finally, we find that

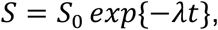

solves the slow pool differential equation.

By fitting an exponential decay to the *t* = 1,2,3,4 minute time points, we obtain the slow fraction *S*_0_ and rate parameter *λ*, which is related to half-life of the slow pool

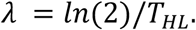

The fast fraction *F*_0_ = *m*_0_ − *S*_0_ and the rate parameter *β* = *λ S*_0_/*F*_0_.

### Estimation of the mRNA decay rate and half-life

The mRNA decay rate and half-life were estimated through the analysis of the decay data, as described in Figure 2 and Supplementary Figure 4. Using the *R* statistics program (85), the data were fitted to a single decay exponential model, represented by the equation, *S* = *S*_0_ *exp*{−*λt*}. The plotting of the data was done using Prism 8 (GraphPad).

### Statistics and Reproducibility

Statistical details can be found in the figure legends. Data were plotted using Prism 8 (GraphPad), correlations were determined using linear regression and the goodness of the fit (r^2^) were reported in the figures. All the experiments presented have been reproduced at least twice with the same results.

## Results

### A richness in the first codons increases protein expression

As it was shown in our previous global study performed in *E. coli (13)*, the base composition of the first 18 nucleotides (6 codons) of the coding sequence, which comprise the area covered by the 70S initiation complex, has a strong impact on protein expression. Increase in A and decrease in G in this region show statistical correlation with protein expression. We decided to use the Venus YFP, as it is one of the most versatile and brightest fluorescent proteins (86) to investigate the role of this region in protein expression. First, we changed the 6 codons after the ATG of the YFP to synonymous ones, by either choosing ones with higher AT content for the YFP_A_ construction or by using the best synonymous tail codons according to our previously published codon metric (13) for YFP_M_ construction (**Figure 1A**). Secondly, we searched in *E. coli* genome for a highly-expressed gene (based on proteomic measurement) that is enriched in A in its first 7 codons and selected the enolase gene. We fused its first 21 nucleotides (which we call a “Translation Boosting Tag” (TBT)) to our YFP reporter (**Figure 1A**). Then, starting from the TBT-YFP construct (TBT), we took the reverse methodology *i*.*e*. replacing TBT codons by their synonymous codons containing the highest G fraction (TBT_G_). To maintain near-physiological conditions, we utilized a low-copy plasmid (pMMB67EH plasmid (77)) expressing the YFP gene under the control of an IPTG-inducible pTac promoter, allowing expression by the native *E. coli* RNA polymerase.

**Figure 1:**
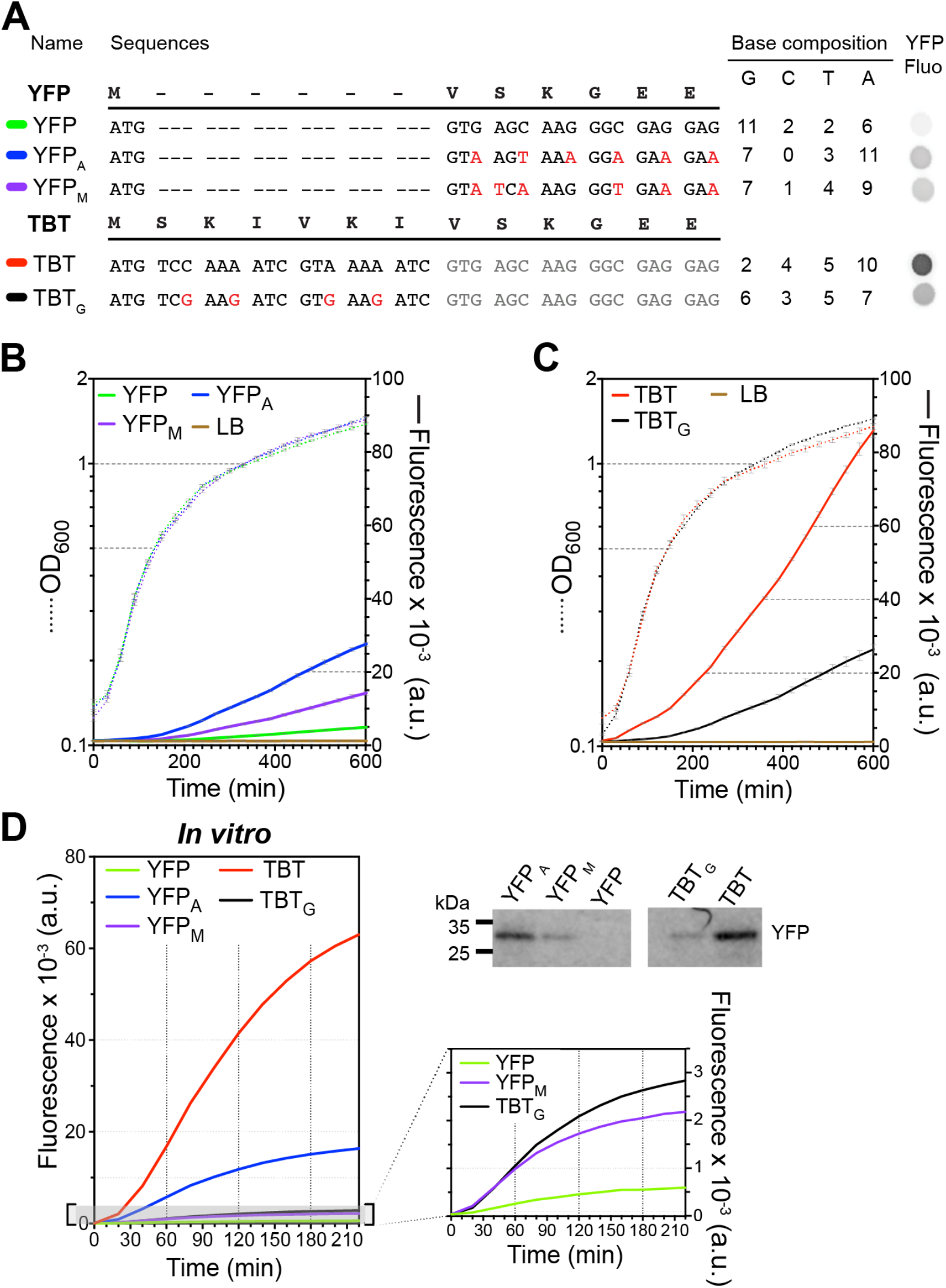
Influence of A-richness in initial codons on protein expression. **A)** Sequences of YFP constructs with synonymous codon replacements in the first 7 codons (YFP_A_, highest A content; YFP_M_, using our model M optimal tail codons (13)). Below, YFP constructs with TBT sequence added at the 5’ end and TBT sequence with replacement by synonymous codons rich in G (TBT_G_). Right, base composition of the constructs and drop assay of different variants, showing the different level of fluorescence *in vivo* on solid medium. **B and C)** *In vitro* measurement of growth and fluorescence of *E. coli* performed in 96-well plate, in LB medium in presence of 1 mM IPTG, **B)** for the YFP and **C)** for the TBT variants. **D)** Fluorescence signal of *in vivo* translation for YFP and TBT variants (PureExpress translation assay with a final concentration of 1.4 µM for each mRNA). Top right, western blots of the final translation reactions revealed with an anti-GFP antibody

The protein production of these YFP variants was first determined by measuring the fluorescence of colonies on agar plates (**Figure 1A, right**). From this result, we can see that all the constructs expressed more than the original YFP and that the TBT construct expresses the most. Then to have a more quantitative and dynamic measurement of the expression, we measured the fluorescence during *E. coli* growth in liquid culture after IPTG induction at the start of the culture (**Figure 1B**). The lag in the fluorescence signal is due to the time necessary to have enough YFP expression to generate a fluorescence signal above the background and to the maturation time of the Venus YFP, which is 17.6 minutes (86). We estimated the protein synthesis rate (defined later one as *dP*/*dt*, in the mathematical model) by measuring the increase per minute of the fluorescence normalized to the OD_600_ in the early linear part of the expression (**Supplementary Figure S2A**). This measurement will also include the refolding and maturation time for the YFP. These variables are consistent for constructs with the same aa sequences.

For the constructs with different aa sequences, we verified that the protein amounts and fluorescent signals were in good agreement, as assessed by protein estimation on Coomassie-stained SDS-PAGE gels or for constructs with low expression by Western blotting using an anti-6xHis antibody (**Supplementary Figure S3D and S4D**). The protein synthesis rate varies widely across the constructs: 3.6 min^-1^ for YFP, 14.9 min^-1^ for YFP_M_, 34.6 min^-1^ for YFP_A_, 29.9 min^-1^ for TBT_G_ and 73.3 min^-1^ for TBT (**Figure 1B and C and Supplementary Figure S2A**). These results again support that the A abundance in the initial codons being a critical determinant of protein expression level, and that codons enriched in A in this region perform better than the optimal codons identified in the remainder of the codon sequence in our earlier study (13). Limitation in A content in synonymous mutations in this region can be surpassed by adding the TBT sequence, as demonstrated by the observation the TBT variant exhibits the highest expression among all YFP constructs (**Figure 1C and Supplementary Figure S2A**). Further supporting the influence of A bases in this region, synonymous codon replacement with an increase in G content in the TBT_G_ construct significantly reduces expression to a similar level of the YFP_A_ construct. These findings highlight that incorporating the first six codons from a highly expressed *E. coli* gene can enhance the expression productivity of a poorly expressing gene with a low (A minus G) value. Furthermore, the results demonstrate a fourfold reduction in expression for the TBT constructs when A is decreased and G is increased.

After 10 hours of growth, the TBT construct expressed protein at a level close to 3 times higher than the YFP_A_ construct (**Figure 1A and B**). While YFP_A_ has slightly more A in the first six codons than TBT (11 *vs*. 10), it also has much more G (7 *vs*. 2). Therefore, the lower protein expression from the YFP_A_ construct could be caused by its greatly increased G content in this region. Although the difference in the amino acid sequence could also potentially influence the difference in the translation efficiency of this construct compared to the TBT construct. In experiments changing the amino acid sequence in the coding region, it is challenging to distinguish the influence of mRNA sequence variations *versus* protein sequence variations on translation efficiency. Several experimental methods have been developed to address this issue (45,87,88) including most impressively *in vitro* tRNA mischarging experiments that support an influence of Lys on the processivity of translation at the start of the coding sequence (45). This elegant research addressed amino-acid effects in a single sequence motif under conditions in which the translation elongation rate is at least an order-of-magnitude slower than it is *in vivo*. To explore whether equivalent effects occur in the TBT and YFP_A_ sequences, we explored the effects of a set of joint variations in their mRNA base composition and amino acid sequences (**Supplementary Figures S2A, S3A-D and S4A-D**). These data, which are described in detail in the ***Supplementary Information*** for this manuscript, support base-content effects being stronger and more predictable in these sequences than amino acid effects. They demonstrate that reducing A content in the first six codons consistently reduces protein expression to at least some extent, while not providing direct evidence supporting an amino-acid effect. The results (**Supplementary Figures S3A-D)** also show that substitution of GAG (Glu) with AAA (Lys) at position 6 (YFP_E6K_) or at positions 6 and 7 (YFP_E6-7K_) produced similar ≈2.3-fold increases in protein synthesis rates (8.4 and 8.3 min−1, respectively compared with 3.6 min−1 for wild-type YFP). The fact that the identical substitution at codon 7 alone did not change expression is consistent with previous reports that the effect of A-enrichment on protein expression is restricted to the first six codons. However, further investigation will be needed to rigorously dissect their relative influence of codon variations *vs*. amino acid variations in these sequences and others.

### The A richness of the first codons improve in vitro protein expression

*In vitro* translation assays (IVTA) programmed with purified mRNA of the different constructs were conducted (**Figure 1D and Supplementary Figure S2B, S3C and S4C**) in minimal translation assay (PURE express from NEB). The fluorescence signal of the YFP was monitored and western blots of the final reaction revelated with anti-GFP antibody confirmed that the fluorescence signal correlates with the YFP amount (**Figure 1D**). Even if we cannot directly link the *in vivo* and *in vitro* kinetics because the ratios of mRNA/ribosome are different, it is noticeable to see that the YFP has a similar protein synthesis rate in both conditions (3.6 min^-1^), whereas TBT expression rate increases by more than 6 times *in vitro* (493.7 min^-1^ *in vitro* vs 73.3 min^-1^ *in vivo*) (**Supplementary Figure S2B**). Overall, all the constructs show a gain of expression when the frequency of A increases, this confirms that the observed effect of the first codons mostly influences mRNA translation *in vivo* and *in vitro*. Nevertheless, some constructs perform differently compared to *in vivo*. YFP_StoK_ is more expressed *in vitro* than the YFP with a rate of 9 min^-1^ vs 3.4 min^-1^. On the other hand, TBT_G_ is less expressed *in vitro* than *in vivo*, making the difference with the TBT construct even more pronounced. The translation of these mRNAs is likely modulated by factors that are present *in vivo* but not in the IVTA, which is composed of the minimal set of translation factors and ribosomes (89).

### Steady state mRNA levels of each construct correlate with their protein expression level

We determined the mRNA levels in a steady state for various YFP constructs through northern blot analysis at different time points after induction with IPTG (0, 10, 60 and 120 minutes) (**Figure 2 and Supplementary Figure S3D**). The northern blot utilizes a DNA probe that hybridizes with the 3’UTR of the mRNA. The TBT construct exhibits the highest mRNA concentration, while YFP has the lowest as they do for protein production. Overall, for all constructs, the mRNA concentration correlates with the produced protein levels. The same trend is observed for the YFP and TBT amino acid variants (**Supplementary Figure S3D and S4D)**. Given that all constructs were expressed using the same pTac promoter, we hypothesize that the variation in mRNA levels is caused by RNA degradation rather than transcription. To assess mRNA stability, we inhibited transcription with rifampicin after 10 minutes of induction by IPTG and determined by northern blot the mRNA concentration at 1-minute intervals (**Figure 2**).

**Figure 2:**
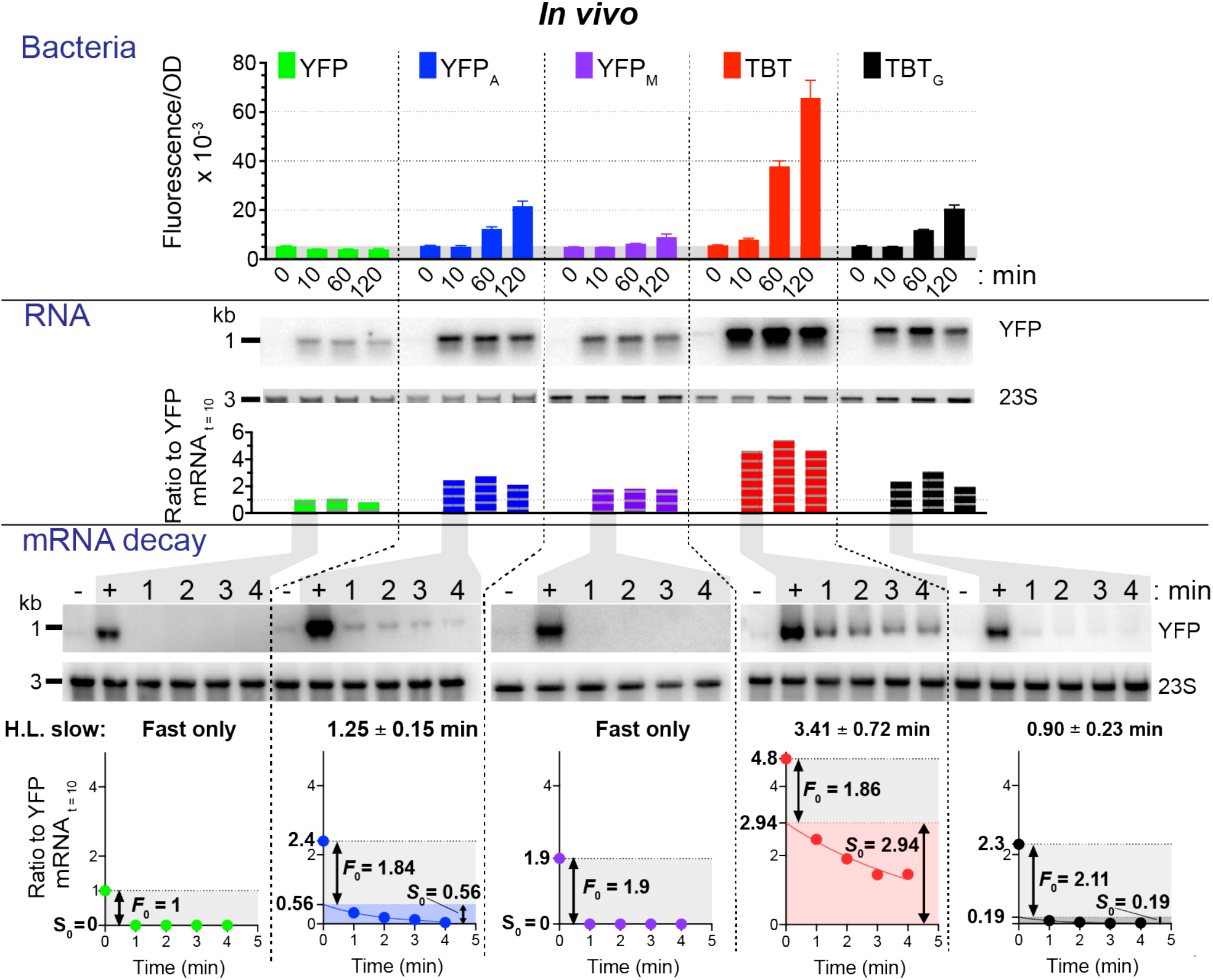
Influence of A-richness on mRNA stability. Fluorescence levels of culture of the different variants collected 10 min, 60 min and 120 min after induction at OD_600_=0.5. Below, northern blotting of total RNA from the corresponding cultures using a DNA probe that hybridizes with the 3’UTR of the *yfp* mRNA. The level of 23S rRNA is showed beneath the northern blot. In the lower section, a mRNA stability test is presented. Northern blot of total RNA isolated from the same strains using the same induction procedure as above. Transcription was halted by rifampicin addition, and samples were collected before induction (-), after 10 min induction (+) and at 1, 2, 3 and 4 min after rifampicin addition. Below, quantification of the northern blot signal normalized to the *yfp* mRNA signal at 10 min after induction along with curve fitting of the decay to an exponential decay for the slow decay part of the kinetic. The proportion of Fast decay (*F*_0_) and Slow decay (*S*_0_) is indicated and the rate of the *S*_0_ decay is indicated above the graph.

After just 1 minute of transcription inhibition, the full-length messenger RNA for the YFP and YFP_M_ constructs was undetectable, but for YFP_A_ and the two TBT constructs a population of more stable mRNA was detectable. The signals before and after rifampicin treatment clearly suggests two populations: one with a fast decay (much less that 1 min) and for some constructs (YFP_A_ and the two TBT), a second one with a slower rate for which we can calculate the half-life (1.25 min for YFP_A_, 0.9 min for TBT_G_ and 3.41 min for TBT). The signal was quantified and normalized relative to the YFP construct’s mRNA concentration before rifampicin treatment. For the constructs showing a slow decay population, the fraction of the fast decay population (*F*_0_) was extrapolated by fitting the slow decay to an exponential decay and taking the intercept with the y axis as the proportion of slow decay (*S*_0_); the *F*_0_ being the difference between the steady-state mRNA concentration and the *S*_0_ fraction. The YFP aa variants (**Supplementary Figure S3D**) also show only a fast decay, but the TBT aa variants show the two decay rates (**Supplementary Figure S4D**).

### Translation by the ribosome is the major determinant of mRNA stability

We looked for correlations between the experimentally determined rates and populations. First, we looked at correlation between the protein expression determined by the YFP fluorescence *in vivo* and the fast and slow mRNA populations, *F*_0_ and *S*_0_ (**Figure 3A**). *F*_0_ is not correlated with fluorescence (r^2^ = 0.05), suggesting that the *F*_0_ population of mRNA is not translated by the ribosome. On the other hand, the *S*_0_ magnitude shows a clear correlation with protein expression rate (r^2^ = 0.82) suggesting that this population of mRNA is the productive one. Secondly, correlation between the expression *in vivo* as monitored by the fluorescence intensity and the steady state mRNA concentration (10 min after induction) shows a linear correlation (r^2^ = 0.96) between the two variables (**Figure 3B**) confirming that the amount of mRNA and translation are linked. Unexpectedly, *in vitro* expression of the constructs in the IVTA with a fixed concentration of each mRNA (1.4 µM) also correlates (r^2^ = 0.82) with the *in vivo* steady state mRNA (**Figure 3B**). Since the only common parameter is the translation of the mRNA by the ribosome, it clearly demonstrates that the slow versus fast decay is dependent on the translation efficiency.

**Figure 3:**
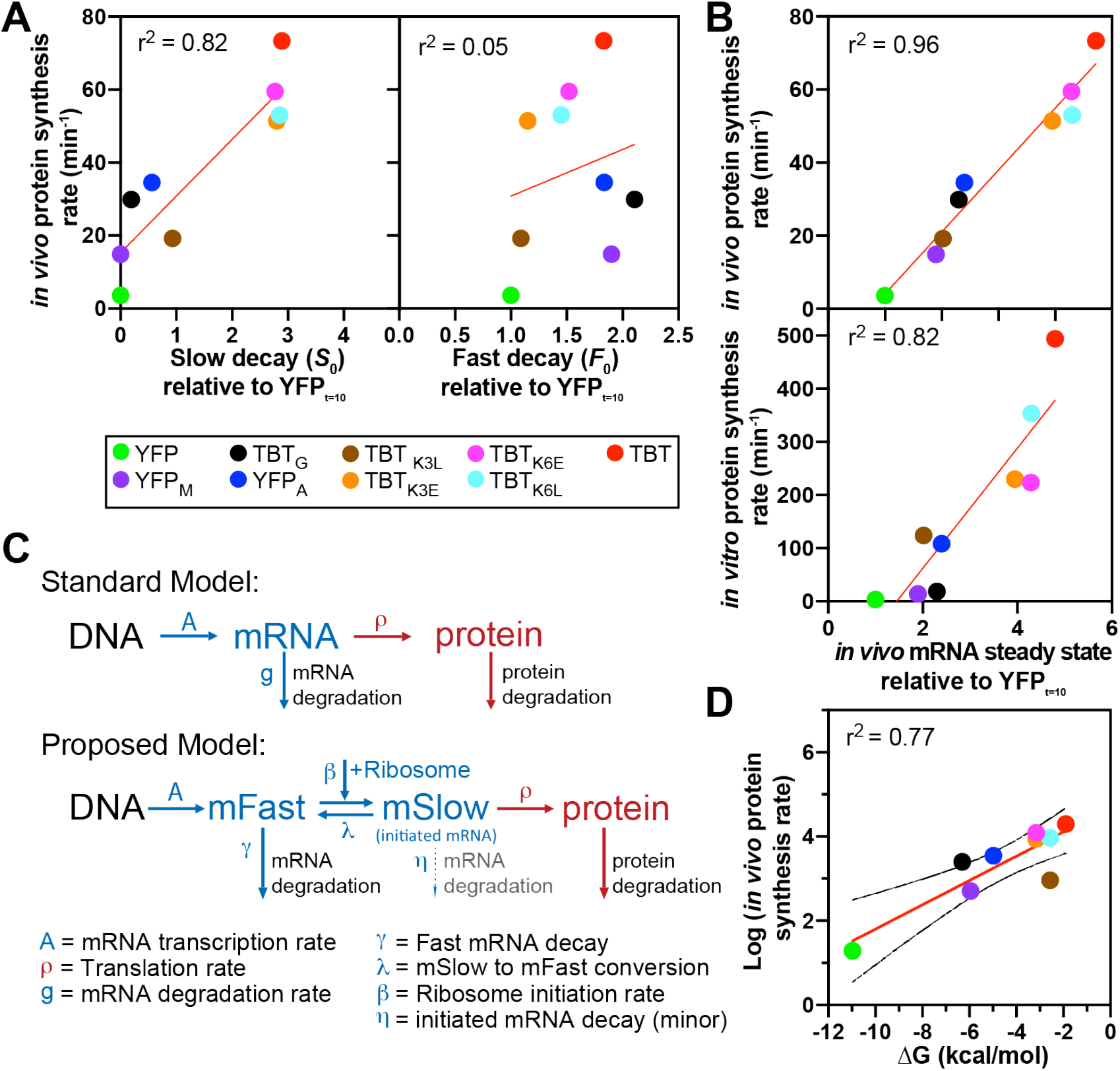
The change in mRNA stability is governed by the ribosome. **A**) The *in vivo* fluorescence increase correlated with the *S*_0_ or *F*_0_ populations (normalized on the *yfp* mRNA signal at 10 min after induction). **B**) Correlation of the *in vivo* or *in vitro* fluorescence increase with the *yfp* mRNA signal at 10 min after induction (normalized on the *yfp* mRNA signal). **C**) Standard model of mRNA decay and our proposed model including the various variables used to construct the model. **D**) Correlation of the logarithm of the *in vivo* fluorescence increase with the energy of folding (ΔG) of the first 21 nucleotides calculated using the BindOligoNet G_5_ model (73). In **A, B, D** the red line is the linear regression, the r^2^ of the goodness of the fit is reported in the right left corner of each graph. In **D** the doted lines show the error.

### Mathematical model that explains the two-mRNA decay rates

In a standard “Central Dogma” model (**Figure 3C**), mRNA (*m*) is transcribed and degraded, and used as a template for protein production (*dP*/*dt*), which yields a simple dynamical model:

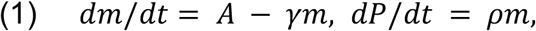

with transcription rate *A*, mRNA degradation rate *γ* and translation rate *ρ*. In steady state,

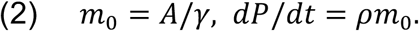

This simple model predicts that the *in vivo* protein production rate *dP*/*dt* increases proportional to the steady-state mRNA level*m*_0_; however, we notably observe in **Figure 3B** that the protein production rate instead goes to zero at a finite mRNA amount,

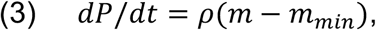

while the simple model has no such *m*_*min*_ offset.

In the decay experiments, when rifampicin shuts off mRNA transcription, *A* → 0. The standard Central Dogma model solution predicts a simple exponential mRNA decay *m*(*t*) = *m*_0_*exp*(−*γt*). However, the experiments instead show a biphasic decay, with a fast portion *F*_0_ decaying within the first minute, and a slow portion *S*_0_ decaying with half-lives ranging from 1 to 4 minutes (**Figure 2, 3A and Supplementary Figure S4D**). To mathematically describe the biphasic decay, we propose a new model that contains two categories of mRNA, *m* = *F* + *S*, with Fast *F* and Slow *S* decaying populations,

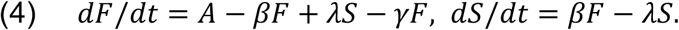

The parameters *β* and *λ* are the rates of conversion between the Fast and Slow populations (**Figure 3C**). We propose that *β* is the 70S ribosome initiation complex formation rate, which is sensitive to the mRNA folding at the beginning of the coding sequence. The steady-state values are:

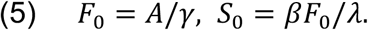

Note that the Fast proportion *F*_0_ is determined by the transcription rate *A* and fast degradation rate *γ*, so *F*_0_ is predicted not to depend on the ribosome-binding rate, which is consistent with **Figure 3A**. However, the Slow component’s proportion *S*_0_ increases with ribosome binding rate *β* and thus with initiation rate.

After rifampicin treatment, *A* → 0, and given a rapid decay rate *γ* ≫ 1/min, we assume the fast portion *F*_0_ has essentially disappeared by the 1 min time point (captured with the delta-function below). This allows us to fit the **Figure 3C** decay data to

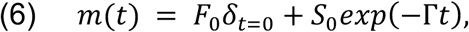

which is essentially a single exponential decay to the *t* = 1,2,3,4 minute time points in that experiment. Extrapolating to *t* = 0 thus gives *S*_0_ and *F*_0_.

Among the various construct sequences, we see that the *in vivo* protein production rate is proportional to *S*_0_, but is independent of *F*_0_, as shown in **Figure 3A**. This finding is consistent with our interpretation that *β* is the 70S ribosome formation rate. Statistical physics suggests *β* is proportional to the Boltzmann likelihood of the ribosome binding site on the mRNA being single-stranded,

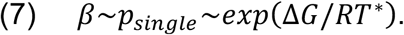

We previously computed the average RNA free energy cost to unfold a portion of an mRNA, finding the four bases contribute unequally (73); with our BindOligoNet model, we predict the free energy cost

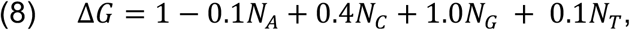

(in kcal/mol) to unfold a region of mRNA. Increasing the number of adenosines *N*_*A*_ and decreasing the number of guanosines *N*_*G*_ has the strongest effect on the initiation rate and protein production. Indeed, we find a Pearson correlation of r = 0.88 (r^2^ = 0.77) between Δ*G* and the logarithm of the protein-production rate in **Figure 3D**, as predicted in the Boltzmann formula above. The thermal fluctuation scale is *RT*=0.615 kcal/mol. We note that the model parameters in Eq (8) for G and A differ by 1.1 kcal/mol = 2 *RT* so their thermal stability is very different, while C and U differ by only 0.3 kcal/mol = 0.5 *RT* which is a relatively less influential. The RNA base pairs GC, AU, and GU permit G and U bases more opportunities to pair than C and A bases, which is why the average energy parameters per base are not equal. Accordingly, the present study concentrates on the effects of A and G; the roles of U and C remain to be investigated in future work. In **Figure 3A**, we see that *F*_0_ across the different constructs is uncorrelated with protein expression, consistent with the steady-state formulas above. Thus *F*_o_ ≈ *m*_*min*_ in Eq. (3).

### Increase of A in the first codon promotes the formation of the 70S IC complex

To understand the molecular mechanism that leads to the expression defect, we compared the TBT and TBT_G_ mRNA abilities to bind the 30S ribosomal subunit using a filter binding assay. The assay demonstrated that the two constructs bind to the 30S subunit with the same affinity (**Figure 4A**). Therefore, the A content in the first codons seems to not affect the interaction between the 30S subunit and the mRNA. However, it is possible that the mRNA is not correctly positioning itself within the mRNA channel of the 30S ribosomal subunit.

**Figure 4:**
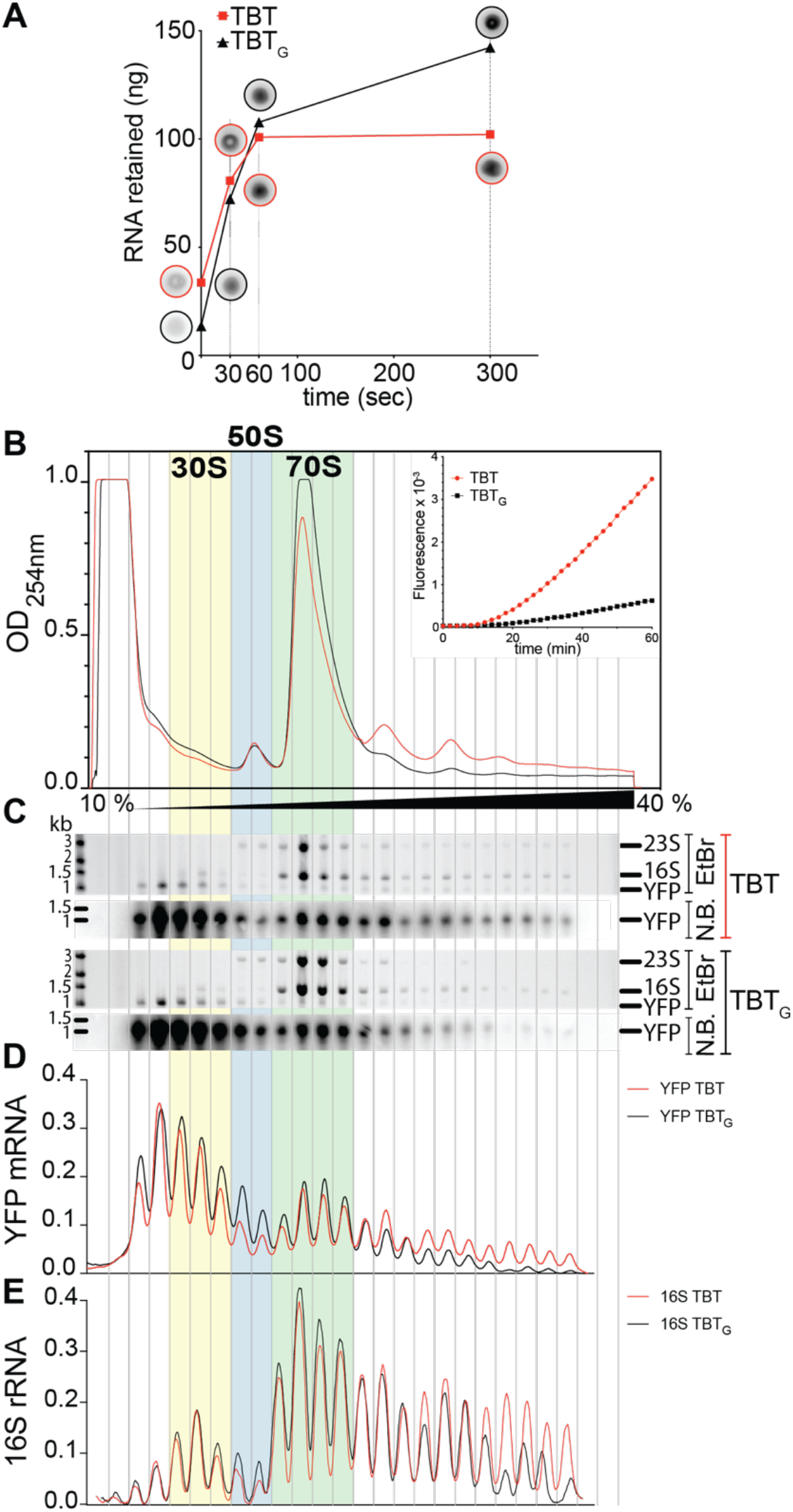
Change in A content for the TBT construct affects the formation of the 70S IC ribosome but not its affinity for the 30S subunit. **A**) Filter binding assay of the P^32^-labeled mRNA (TBT or TBT_G_), quantified after different incubation time with the 30S ribosomal subunit. Image filter after scanning are presented in the circles. **B**) Polysomes separation and fractionation of IVTA programed with the TBT or TBT_G_ mRNAs after 1 h of incubation (the IVTA is presented in the insert). **C**) Ethidium bromide stained gel (EtBr) or northern blot (N.B.) of the total RNA purified from each fraction of the polysome profiles. **D**) Normalized graph of the quantification of the northern blot signal for the TBT or TBT_G_ mRNAs. **E**) Normalized graph of the quantification of the Ethidium bromide stained gel signal for the 16S rRNA.

To gain an overview of the differences during the translation of the two mRNAs, we conducted an IVTA with each mRNA. After 1-hour of reaction, we separated each reaction on sucrose gradient, collected the fractions and extracted the total RNA (**Figure 4B**). We first tried to use the fluorescence of the MGapt aptamer (82,90) that we added in the 3’UTR of our transcript, but the fluorescence signal was too low for detection (data not shown). Therefore, we determined, through northern blot analysis, the location of the mRNA within each profile. Several observations emerged from this experiment. First, the TBT_G_ mRNA produced fewer polysomes, and consequently, less mRNA was found in the polysome fractions (**Figure 4C**), suggesting a reduced entry of ribosomes into the elongation cycle. Secondly, depletion of TBT_G_ mRNA in the polysomes led to an accumulation in the 30S-50S fractions (**Figure 4C and 4D**). This accumulation correlates with an increase in 16S rRNA in the 50S fraction, indicating a trailing of the pre-30S initiation complex within the 50S location. From these observations we conclude that the constructs have a similar affinity for the 30S ribosomal subunit and that the TBT_G_ variant affects the formation of the 70S Initiation Complex (IC) after formation of the 30S pre-IC or the 30S IC.

The depletion of polysomes during the translation of TBT_G_ mRNA, as discussed earlier, indicates a diminished influx of ribosomes into the elongation cycle. This could be induced by reduced functional initiation or by abortive early elongation cycles (45,48-50). Abortive elongation leads to the release of peptidyl-tRNA from the elongation complex, which dissociates during the elongation process. To test this hypothesis, we conducted a peptidyl-tRNA drop-off assay (**Supplementary Figure S5**). IVTA coupled with T7 transcription was performed in the presence of [^35^S]-methionine and DNA templates of *yfp, yfp*_*a*_, *tbt* and *tbt*_*g*_ with T7-promoter. Following precipitation and washing, the samples were separated by electrophoresis. For each sample, half was treated with RNase to identify tRNA and peptidyl-tRNA species. These assays did not reveal any peptidyl-tRNA for any of the tested samples, suggesting abortive translation does not contribute significantly to the protein expression effects observed in our experiments.

### The TBT sequence can generally be used to increase protein production

To assess whether the TBT sequence can enhance the expression of proteins other than YFP, we fused it to the 5’-end of genes encoding proteins with poor expression levels. Genes *ycaQ* (*E. coli*), RSP_2139 (*Rhodobacter sphaeroides*), SCO1897 (*Streptomyces coelicolor*) and SRU_1983 (*Salinibacter ruber*) were previously assessed for their expression in the *E. coli* Bl21 DE3 strain (13). This evaluation was conducted utilizing a pET21 plasmid, placing the expression of the gene under the control of a T7 promoter. The WT constructs exhibited negligible to very low expression levels (13) (**Figure 5A**). Upon introducing the TBT tag fused to the N-terminus of the protein, we observed a significant improvement in the expression of *ycaQ*, SRU_1983 and RSP_2139, as well as an observable gain in the expression of SCO1897 (**Figure 5A**). Interestingly, all tested genes have low A content and high G/C content. These findings underscore the significance of the first codons in achieving higher protein expression and highlight the TBT tag as an accessible and cost-effective option for boosting the expression of certain proteins. They also demonstrate that the A richness effect of the early codons is not dependent on the native polymerase of *E. coli* and can be observed when genes are transcribed with a phage T7 polymerase, which uncouples transcription from translation (5,91).

**Figure 5:**
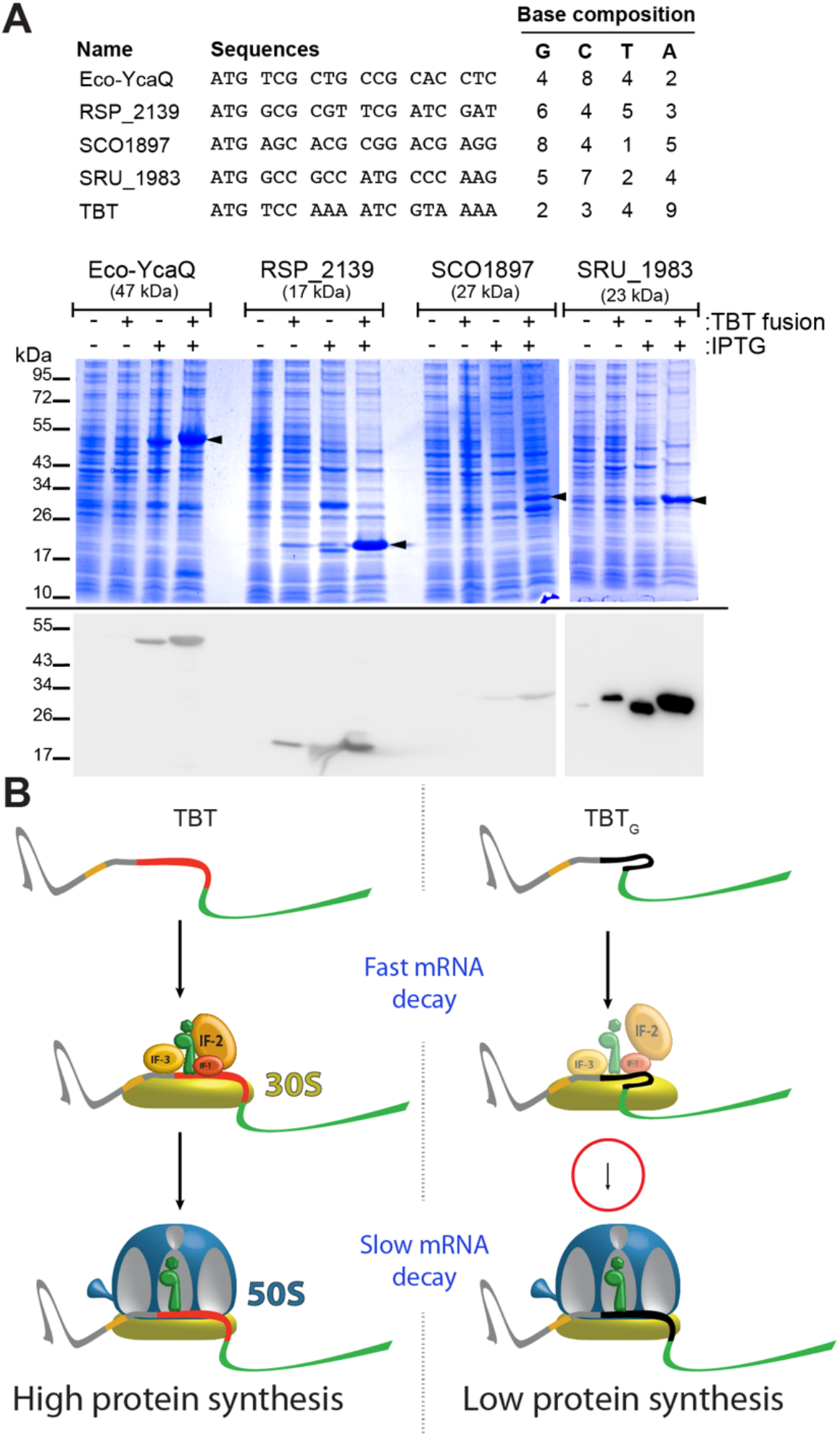
Use of the TBT sequence to improve protein expression and speculative mechanism of its effect on translation initiation and mRNA decay. **A**) Four genes that are poorly expressed in *E. coli* (13) were tested with or without a 5′ TBT fusion. Top: table showing the sequences of the first six codons of each tested gene and the TBT sequence. Bottom: SDS-PAGE showing induction or not of tested proteins (black arrows) with their WT sequence or the sequences with the TBT sequence fused in N-terminal. Protein induction was performed as previously (13) in the *E. coli* Bl21 DE3 strain with 1 mM IPTG for 16 h at 17 °C. Below, Western Blot with anti-his antibodies showing induction of tested proteins. **B**) Model illustrating the impact of A-richness in initial codons, derived from observations of TBT vs. TBT_G_. The TBT mRNA rapidly assembles into a 70S initiation complex (IC), whereas the TBT_G_ mRNA exhibits slower 70S IC formation, getting trapped at the 30S IC formation stage. It remains uncertain whether the kinetic restriction occurs before or after the assembly of initiation factors and the initiator fMet-tRNA^fMet^. The free pool of mRNA and the stalled 30S IC or pre-30S IC with the TBT_G_ construct undergo rapid degradation, while the mRNAs engaged in the 70S IC are stabilized, leading to a slower decay.

## Discussion

In this study, we present results supporting a model (**Figure 5B**) in which the base composition of the first six codons controls both the translation efficiency and stability of mRNAs in *E. coli*. This effect is mediated by the formation of the 70S IC, which is promoted by early codons enriched in A and depleted in G. The A richness of the mRNA does not seem to affect its binding affinity for the 30S subunit, but it does influence the formation of the 70S IC. Free mRNAs and those bound to the 30S but not matured into the mRNA 70S IC complex are non-productive, forming a population of rapidly decaying mRNA. This model implies that successful initiation of translation by the ribosome is a major factor dictating mRNA stability, a proposal that aligns with the established understanding of this step as a pivotal checkpoint in mRNA selection (92). This proposal provides a more concrete explanation of the mechanism outlined by previous study suggesting ribosomal protection of mRNA (93-95).

The sequence of the TBT motif (MSKIVKI), corresponding to the initial codons of the enolase gene, includes a motif described by Verma *et al*., specifically K/N-Y/I at positions 3 and 4 (45). Our selection of this sequence in our study predates their publication; thus, the independent identification of this motif underscores its significance. In their study, the authors suggest that the aa motif helps prevent stalling and abortive early elongation events. Although our analysis does not possess the extensive power of the deep eGFP library used in their research and examines only a limited number of constructs, we clearly observe that the A vs. G ratio within the TBT sequence modulates expression. The TBT construct produces 3.4 times more YFP than the TGT_G_ construct *in vivo* (**Figure 1C**) and 20 times more *in vitro* (**Figure 1D**). While we cannot exclude the possibility that that some of the tested sequences alter the level of abortive translation (45,48,96), none of the tested constructs (YFP, YFP_A_, TBT, and TBT_G_) demonstrated any evidence of peptidyl drop-off when evaluated using a standard *in vitro* assay (**Supplementary Figure S5**). This difference relative to an earlier study with a similar focus (45) suggest that multiple factors influence initiation and early translation events, collectively mediating the overall expression levels of a transcript.

We built a mathematical model (**Figure 3C**), that describes the biphasic mRNA decay observed in our experiments (**Figure 2** and **Supplementary Figure S4D**) by postulating two populations of mRNA. Protein production level is proportional to the slowly decaying mRNA population, while the fast-decaying mRNA population is uncorrelated with protein level (**Figure 3A**). Our model differs from the usual “translation efficiency” assumption (Eq 1) of proportionality between protein rate and total mRNA, instead invoking a substantial population of unproductive mRNA (Eq 3). A Boltzmann factor with an unfolding free energy given by Eqs 7-8 predicts the ribosome initiation rate (**Figure 3D**), explaining why A-rich and G-poor sequences are translated at higher rates. Our model predicts that the initiating ribosome is the pivot that switches the fast-decaying mRNA population toward the slow-decaying population. This model is consistent with a recent publication that demonstrates the primary determinant of mRNA stability in yeast is the initiation of mRNA translation by the ribosome (97).

This model of mRNA degradation should drive directional decay of the mRNA from the 5’ to the 3’, despite the absence of a 5’ to 3’ processive RNase in *E. coli*. The polarity, as previously described in the literature (98,99), may arise from the accessibility and phosphorylation state of the 5’-end of the mRNA for the main *E. coli* RNase, RNase E (100-102). The polarity of decay is then controlled by the scanning ability of the RNase E (103). This model aligns with the ribosomal mRNA protection model (103), but it suggests that interaction with the 30S subunit alone does not provide protection against mRNA degradation but instead controls a competition between degradation and translation initiation. Future research should aim to characterize the molecular conformation and interactions of the abortive 30S IC and how it funnels mRNA to degradation enzymes.

Our model offers an alternative explanation to the “translational ramp” theory, which proposes that enrichment of rare codons near the start of coding sequences slows early elongation to increase translation speed later on. That hypothesis implies that rare codons are translated more slowly, a point that remains highly debated (see Introduction), and which several studies, including ours, have not supported (11,13,15,16,21-33). Here we show that A-enrichment and G-depletion in the first six codons, features commonly associated with rare codons, promote efficient ribosome initiation. Thus, the prevalence of rare codons at the start of ORFs may reflect nucleotide composition that favors initiation (A-rich/G-poor), rather than an evolutionary mechanism to impose a slow-to-fast elongation ramp.

In conclusion, we propose that the ribosome-docking region in an mRNA and specifically the first six encoding codons play a pivotal role in orchestrating a dynamic competition between mRNA decay and translation initiation. Initiation of mRNA decay primarily upstream (5’) of translating ribosomes is a mechanism that reconciles disparate observations (13,15,17,21,45,46,63,104). This paradigm optimizes metabolic efficiency, by ensuring that mRNA decay is initiated upstream of translating ribosomes rather than downstream. This regulatory mechanism will prevent the wasteful expenditure of energy involved in the production and subsequent degradation of partially synthesized proteins.

The research reported in this paper also establishes a cost-efficient and straightforward strategy to enhance protein expression in *E. coli* by fusing target proteins after the Translation-Boosting Tag (TBT) sequence. We showed that this tag can boost the expression of proteins transcribed either by the native *E. coli* RNA polymerase or by the phage T7 polymerase. This tag also enhances protein expression in cell-free *in vitro* translation systems as we show for the YFP (TBT, **Figure 1D**).

## Data availability

The data underlying this article are available in the article and in its online supplementary material.

## Acknowledgement

We thank Farès Ousalem for sharing some of his purified 70S ribosomes.

## Author contributions

AL and LM with the help of SN, TO and CB performed the experiments. AL, LM and GB analyzed the data. IW, JFH, DPA and GB built the mathematical model. AL, LM and GB prepared the figures. GB designed the research program. AL, LM, DPA, JFH and GB wrote the manuscript in consultation with the other authors.

