## Supplementary information for "Base composition at the start of the coding sequence controls the balance between translation initiation and mRNA degradation in *E. coli*"

### *for*

### Supplementary Text

***Evaluation of a set of joint variations in mRNA base composition and amino acid sequence at the start of the protein-coding sequence.*** The presence of positively charged amino acids at the beginning of the coding sequence correlates with increased mRNA translation in bacteria (1) and yeast (2). In *Bacillus subtilis*, a Lys is found at position 2 in 20% of the genes (3). Other studies have also identified Lys at position 3 as being correlated with translation efficiency in other bacteria (4,5). However, the codons for lysine are strongly A-biased (AAA and AAG), meaning these correlations could potentially derive from nucleotide-sequence effects rather than amino acid effects. However, it is fundamentally challenging to deconvolute the effects of amino acid *vs.* nucleotide variations on mRNA translation because amino acid identity cannot be changed without changing nucleotide sequence except in specialized experiments exploiting nonsense suppression effects *in vivo* or tRNA mischarging *in vitro*. Verma *et al*. used elegant tRNA mischarging experiments to demonstrate that charging the same tRNA with Lys *vs.* Val at position 3 increases processivity during *in vitro* translation (1), providing a clear example of amino acid identity influencing mRNA translation albeit under *in vitro* conditions. Given that our TBT construct has Lys residues at both positions 3 and 6, while the $YFP_A$ construct has just a single Lys residue at position 4 (**Figure 1A**), the substantially higher expression we observed for the TBT construct could be influenced by the differences in its N-terminal amino acid sequence in addition to or instead of the differences in its nucleotide base composition in this region. To try to gain insight into their relative influence on the expression of the TBT *vs.* $YFP_A$ constructs, we conducted a series of experiments comparing the effects of changing amino acid identity or exchanging amino acids at positions in the first six codons.

We first substituted the GAG codon for Glu with the AAA codon for Lys just at position 6 or at both positions 6 and 7 in YFP (**Supplementary Figures S2A and S3A-B**). The single-mutant $YFP_{E6K}$ construct and the double-mutant $YFP_{E6\text{-}7K}$ construct showed very similar ~2.3-fold increases in protein synthesis rates (8.4 and 8.3 $min^{-1}$, respectively) compared to YFP (3.6 $min^{-1}$). The modest gain in expression efficiency in the single-mutant $YFP_{E6K}$ construct could be attributable either to the substitution of Lys for Glu or the substitution of two As for two Gs in the sixth codon or potentially influenced by both factors. The failure of exactly the same substitutions at the seventh codon to change expression level is consistent with previous reports that the influence of A enrichment on protein expression is limited to the first 6 codons (6,7).

To test the influence of having a Lys residue specifically at position 3 in the coding sequence, we swapped the Ser codon at position 3 and the Lys codon at position 4 in YFP to change the aa sequence at the N-terminus of the protein without altering the nucleotide base composition in this region (**Supplementary Figures S2A and S3A-B**). The protein synthesis rate of the resulting $YFP_{StoK}$ with Lys at position 3 was ~2.1-fold lower than the parental unswapped YFP construct (1.7 *vs.* 3.6 $min^{-1}$), showing that, at least in the context of the N-terminal amino acid sequence in YFP, having a lysine at position 3 does not increase protein expression when the nucleotide base content in this region remains constant.

To further evaluate the role of Lys residues at the first six positions in the coding sequence, we made the same pair of codon and aa substitutions (GAA for Glu or TTA for Leu) at each of the two Lys residues in this region in the TBT sequence, which are located at positions 3 and 6 and both encoded by the AAA codon (**Supplementary Figure S2A and S4A-B**). The Lys-to-Glu substitutions at both sites produce similar 21-32% reductions in protein synthesis rate (51.4 $min^{-1}$ for $TBT_{K3E}$ and 59.5 $min^{-1}$ for $TBT_{K6E}$ *vs.* 75.3 $min^{-1}$ for TBT). While these effects go in the same direction, they are smaller in magnitude than the ~57% reduction observed when substituting the synonyous GAG codon for Glu for the AAA codon for Lys at position 6 in YFP (**Supplementary Figures S2A and S3A-B**), which doubles the number of A-to-G substitutions used to replace lysine with the same amino acid. These observations reinforce the conclusion that Lys at position 3 does not have a unique influence on protein expression level compared to Lys at other positions in the first six codons, at least not in the context of the N-terminal aa sequence of TBT. They furthermore provide another example of the consistency with which reducing A content in the first six codons in the same sequence context tends to reduce protein expression. The very modest effects produced by these Lys-to-Glu substitutions in TBT could again be influenced by the change in amino acid identity in addition to or instead of the change in nucleotide base composition. Nonetheless, our results tend to weigh against a specific stimulation of protein expression by Lys in the context of our sequences. If the primary effect of these substitutions on translation level were due stereochemical interactions in the peptidyl transferase center or nascent-peptide exit tunnel of the ribosome, which would presumably be responsible for mediating amino acid effects (1,8-10), their influence would be expected to depend on protein sequence position and context. However, the observed effects of Lys-to-Glu substitution on protein expression in our experiments do not.

In contrast to the very similar impact of the substitutions in the $TBT_{K3E}$ and $TBT_{K6E}$ constructs, the substitutions in the $TBT_{K3L}$ and $TBT_{K6L}$ constructs have significantly different effects on protein expression depending on the position (**Supplementary Figure S2A and S4A-B**). Substituting the TTA codon for Leu to replace the AAA codon for Lys at position 6 in TBT produces an ~30% reduction in protein synthesis rate (53.0 $min^{-1}$ for $TBT_{K6L}$ *vs.* 75.3 $min^{-1}$ for TBT), a very similar reduction to that observed when the GAA codon for Glu is introduced at either that position in the $TBT_{K6E}$ construct or at position 3 in the $TBT_{K3E}$ construct. However, a substantially larger ~74% reduction in protein synthesis rate is observed when the same substitution of the TTA codon for Leu is used to replace the AAA codon for Lys at position 3 in TBT (19.3 $min^{-1}$ for $TBT_{K3L}$ *vs.* 75.3 $min^{-1}$ for TBT). Notably, protein expression from the $TBT_{K3L}$ construct is somewhat lower than that from the $TBT_G$ construct even though the former has 8 As and only 2 Gs in its first six codons *versus* 7 As and 6 Gs in the latter. The strong position/context-dependence of the Lys-to-Leu substitutions in TBT, as well as that of the codon swap in the $YFP_{StoK}$ construct reported above, suggests some specific stereochemical interactions contribute to the observed variations in protein expression efficiency, although these interactions could potentially involve either the first six amino acids in the nascent protein or the corresponding codons in the mRNA.

Importantly, in all of the amino acid and codon substitution experiments reported here, reducing A content in the first six codons in a given sequence context consistently reduces protein expression at least to some extent. While our results do not rule out amino acid content in this region influencing protein expression, they do not provide direct evidence for such an effect. In contrast, they provide extensive evidence that increasing A content or decreasing G content in the first six codons increases protein expression, although the magnitude of these effects varies depending on sequence context and position.

### Figure S1

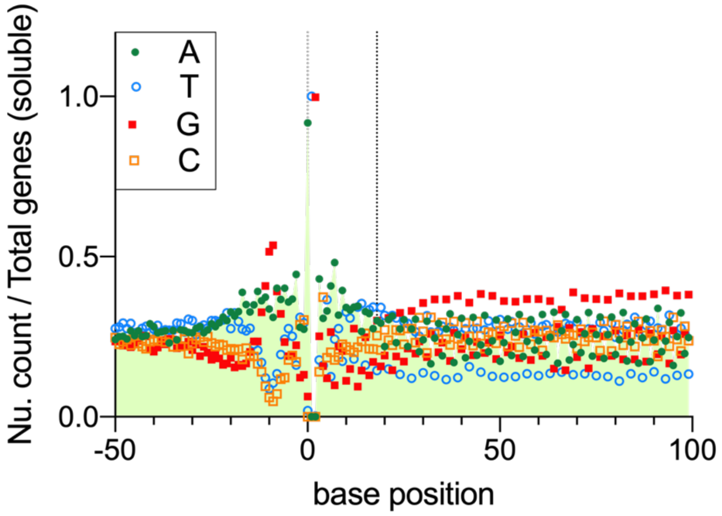


**Figure S1:** ***Base composition of the 5'-end of the coding sequences for soluble proteins in E. coli genome.*** The base composition at each position (0 is the A of ATG) was counted for each *E. coli* gene encoding soluble protein and was divided by the number of counted genes. The dashed line shows the end of the 6 first codons.

### Figure S2

A

B

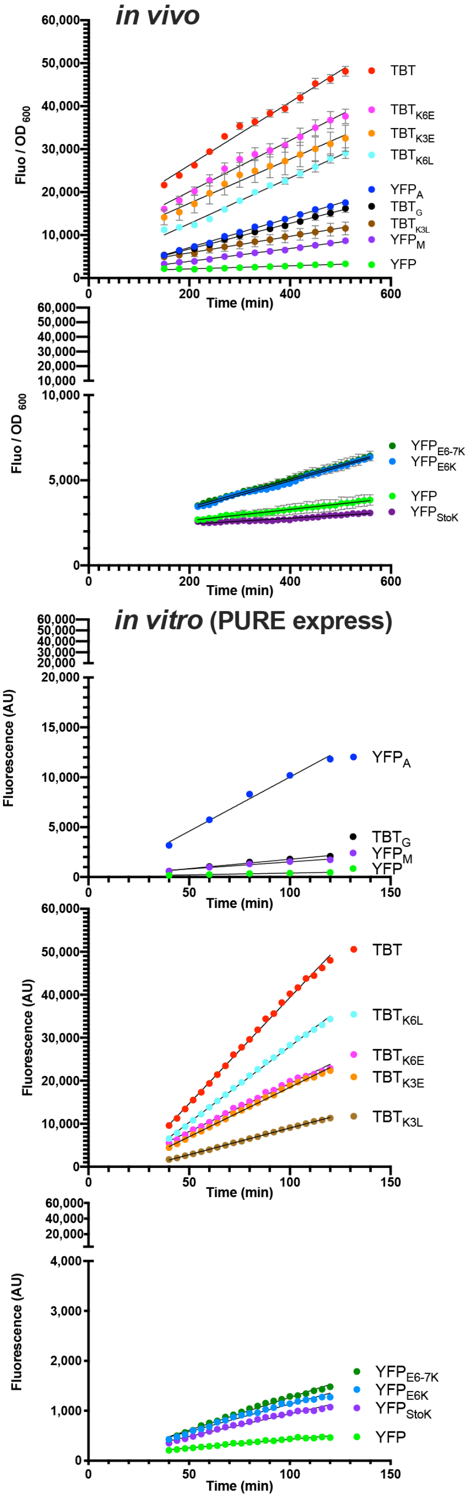


***In vivo*** **Fluo/OD (150 to 510 min)**

| | Slope (min$^{-1}$) | Std. Error |
|---|---|---|
| YFP | 3.6 | 0.1599 |
| $YFP_A$ | 34.6 | 0.4377 |
| $YFP_M$ | 14.9 | 0.428 |
| $TBT_G$ | 29.9 | 0.7989 |
| TBT | 73.3 | 1.681 |
| $TBT_{K3L}$ | 19.3 | 1.42 |
| $TBT_{K3E}$ | 51.4 | 3.316 |
| $TBT_{K6E}$ | 59.5 | 2.156 |
| $TBT_{K6L}$ | 53.0 | 1.211 |

***In vivo*** **Fluo/OD (216 to 560 min) not used in the model**

| | Slope (min$^{-1}$) | Std. Error |
|---|---|---|
| YFP | 3.3 | 0.2309 |
| $YFP_{E6K}$ | 8.4 | 0.1036 |
| $YFP_{E6-7K}$ | 8.3 | 0.1943 |
| $YFP_{StoK}$ | 1.7 | 0.1021 |

***In vitro*** **(40 to 120 min)**

| | Slope (min$^{-1}$) | Std. Error |
|---|---|---|
| YFP | 3.6 | 0.2499 |
| $YFP_A$ | 108.6 | 6.356 |
| $YFP_M$ | 14.1 | 1.294 |
| $TBT_G$ | 18.8 | 1.202 |
| TBT | 493.7 | 5.452 |
| $TBT_{K3L}$ | 124 | 0.7606 |
| $TBT_{K3E}$ | 230 | 2.847 |
| $TBT_{K6E}$ | 223.5 | 4.128 |
| $TBT_{K6L}$ | 353.4 | 3.12 |
| **Below not used in the model** | | |
| YFP | 3.4 | 0.1235 |
| $YFP_{E6K}$ | 11 | 0.3348 |
| $YFP_{E6-7K}$ | 13.1 | 0.2888 |
| $YFP_{StoK}$ | 9.1 | 0.2675 |

**Figure S2:** ***Quantification of the protein synthesis rate (dP/dt).*** The protein synthesis rate is the slope in the early linear part of the (fluorescence / $OD_{600}$) ratio for *in vivo* expression or fluorescence for *in vitro* experiments. The slopes and Std. Error were calculated using simple linear regression in Graphpad Prism 8 software.

### Figure S3

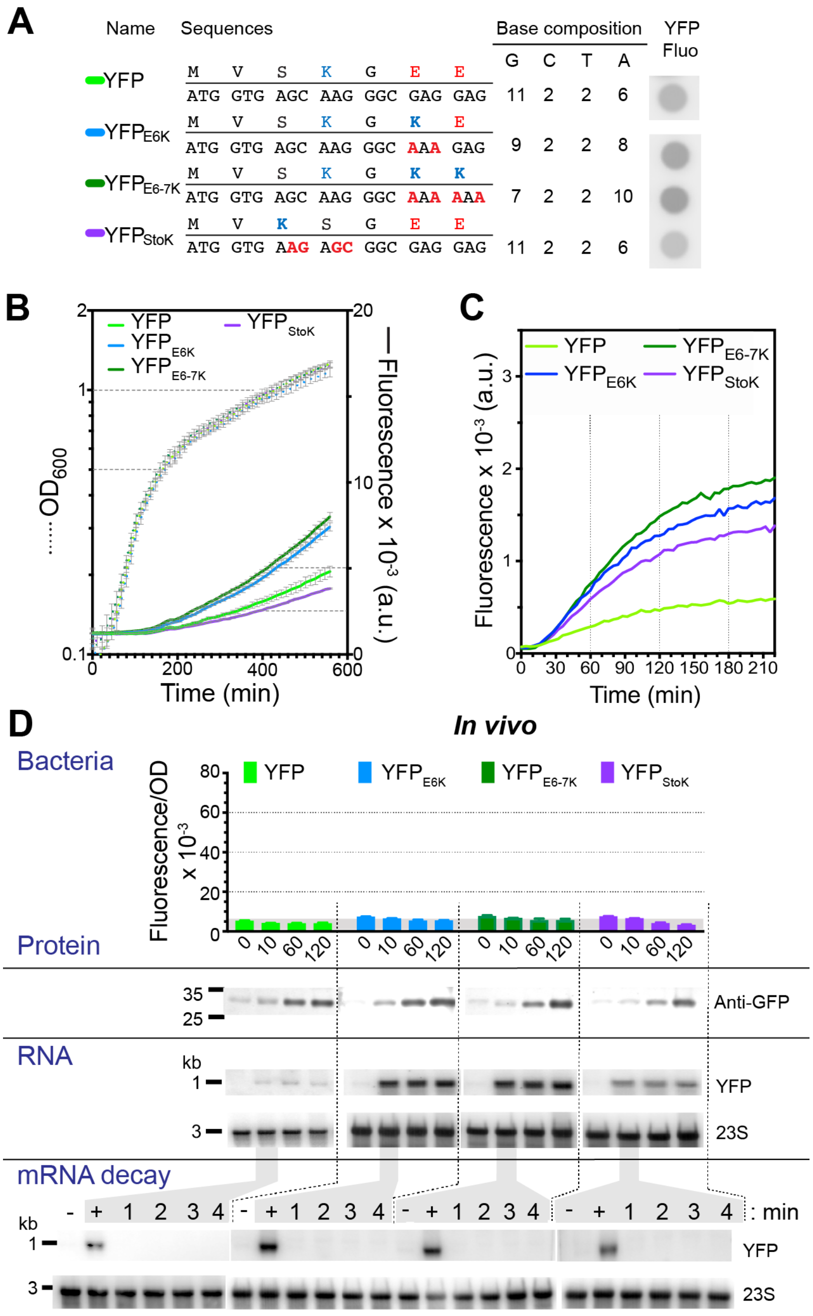


**Figure S3:** ***Influence of the presence of lysines in the first aa of the YFP on its expression and stability of its mRNA.*** **A**) Sequences of YFP constructs with codon replacements for introduction of Lys in the first codons. Right, base composition of the constructs and drop assay of different variants, showing the different level of fluorescence *in vivo* on solid medium. **B**) *In vivo* measurement of growth and fluorescence of *E. coli* performed in 96-well plate, in LB medium in presence of 1 mM IPTG. **C**) Fluorescence signal of *in vitro* translation for YFP lysine variants (PureExpress translation assay with a final concentration of 1.4 μM for each mRNA). **D**) Fluorescence levels of culture of the different variants collected 10 min, 60 min, and 120 min after induction at $OD_{600}$=0.5. Below, western blot of the extracts at the different time points revelated with an anti-YFP antibody. Center, northern blotting of total RNA from the corresponding cultures using a DNA probe that hybridizes with the 3'UTR of the *yfp* mRNA. The level of 23S rRNA is shown beneath the northern blot. In the lower section, mRNA stability test is presented. Northern blot of total RNA isolated from the same strains using the same induction procedure as above. Transcription was halted by rifampicin addition, and samples were collected before induction (-), after 10 min induction (+), and at 1, 2, 3, and 4 min after rifampicin addition. The decay profile for YFP is the same as the one presented in Fig. 2, with the signal intensity adjusted to match that of the YFP lysine variants.

### Figure S4

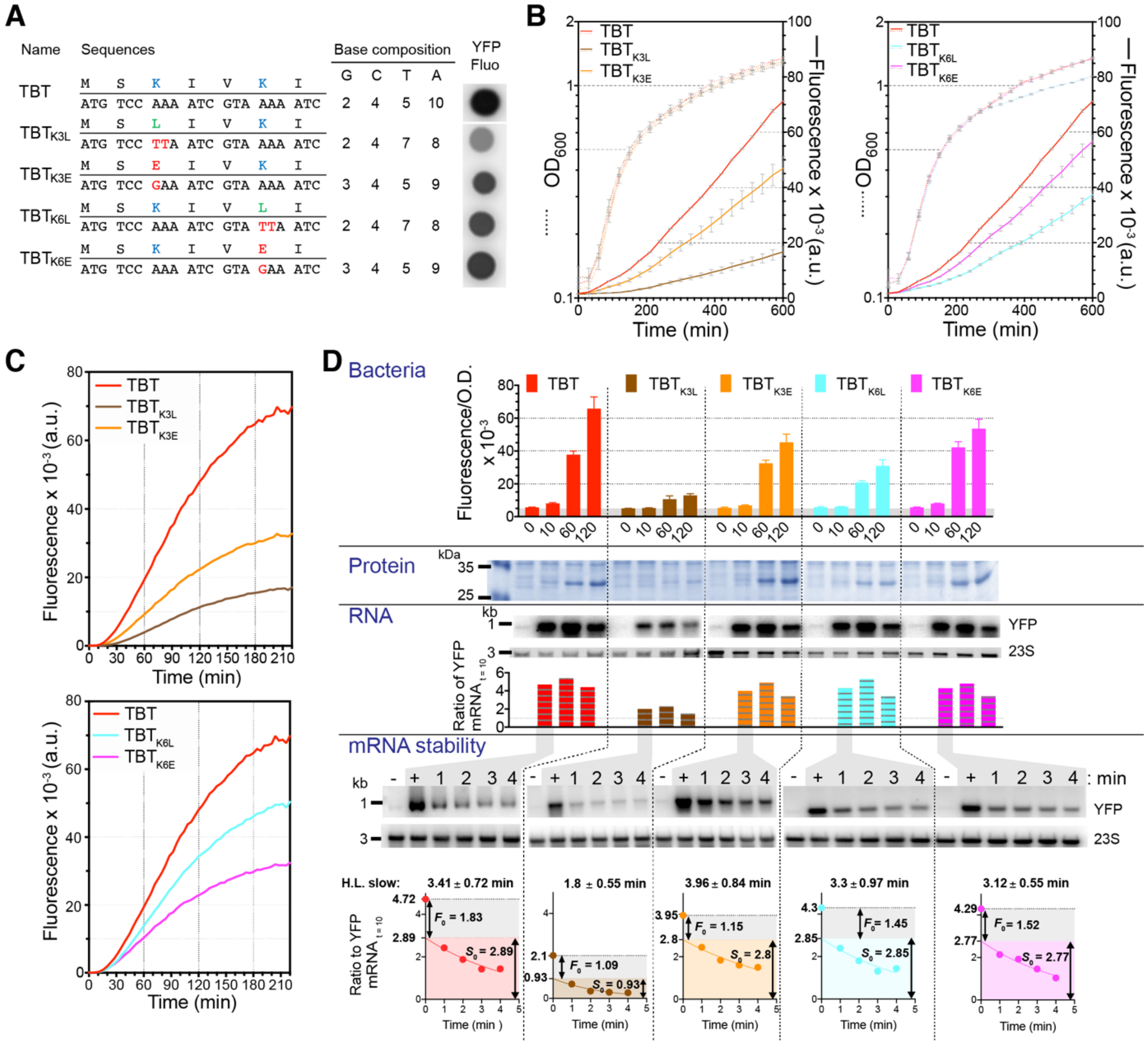


***Figure S4: Influence of the presence of lysines in the first aa of the TBT on its expression and stability of its mRNA.*** **A**) Sequences of TBT constructs with codon replacements for introduction of Lys in the first codons. Right, base composition of the constructs and drop assay of different variants, showing the different level of fluorescence *in vivo* on solid medium. **B**) *In vivo* measurement of growth and fluorescence of *E. coli* performed in 96-well plate, in LB medium in presence of 1 mM IPTG. **C**) Fluorescence signal of *in vitro* translation for YFP lysine variants (PureExpress translation assay with a final concentration of 1.4 µM for each mRNA). **D**) Fluorescence levels of culture of the different variants collected at 10 min, 60 min, and 120 min after induction at $OD_{600}$=0.5 Below, SDS gel of the protein extracts showing a good correlation between fluorescence signal and protein expression. Below, northern blotting of total RNA from the corresponding cultures using a DNA probe that hybridizes with the 3'UTR of the *yfp* mRNA. The level of 23S rRNA is showed beneath the northern blot. In the lower section, a mRNA stability test is presented. Northern blot of total RNA isolated from the same strains using the same induction procedure as above. Transcription was halted by rifampicin addition, and samples were collected before induction (-), after 10 min induction (+), and at 1, 2, 3, and 4 min after rifampicin addition. Below, quantification of the northern blot signal normalized to the *yfp* mRNA signal at 10 min after induction, along with curve fitting of the decay to an exponential decay for the slow decay part of the kinetic. The proportion of fast decay ($F_0$) and slow decay ($S_0$) is indicated, and the rate of the $S_0$ decay is indicated above the graph.

### Figure S5

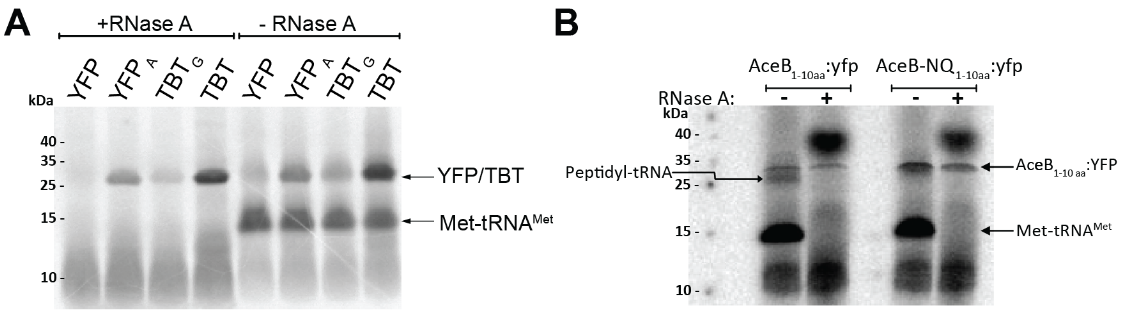


***Figure S5: Analysis of Peptidyl-tRNA Drop-off During Translation.*** **A**) DNA templates promoting the expression of *yfp*, $yfp_a$, *tbt*, and $tbt_g$ were transcribed and translated in the PUREfrex 1 Coupled Transcription/Translation System for 90 minutes in the presence of [$^{35}$S]-Methionine. The products were then separated by electrophoresis on an 11% Wide Range Gel, with or without prior treatment with RNase A. **B**) Control experiment demonstrating peptidyl-tRNA drop-off during the translation of the first codon of the *aceB* gene at the acidic residues D and E at positions 9 and 10. In the variant where the D and E residues are substituted with N and Q, peptidyl-tRNA drop-off does not occur (11).
